# RatHat: A self-targeting printable brain implant system

**DOI:** 10.1101/868422

**Authors:** Leila M. Allen, Maanasa Jayachandran, Tatiana D. Viena, Meifung Su, Timothy A. Allen

**Author notes:** Contributed equally to this work (Leila M. Allen, Maanasa Jayachandran). Author Contributions: LMA, MJ, MS, BLM, and TAA designed research; LMA, MJ, MS, and TDV performed research; LMA, MJ, TDV and TAA analyzed data; LMA, MJ, and TAA wrote the paper. Corresponding Author: Timothy A. Allen, PhD, Department of Psychology, Florida International University, 11200 SW 8^th^ Street, Miami, FL, 33199, Website: http://allenlab.fiu.edu/, Twitter: @AllenNeuroLab.

## Abstract

There has not been a major change in how neuroscientists approach stereotaxic methods in decades. Here we present a new stereotaxic method that improves on traditional approaches by reducing costs, training, surgical time, and aiding repeatability. The RatHat brain implantation system is a 3D printable stereotaxic device for rats that is fabricated prior to surgery and fits to the shape of the skull. RatHat builds are directly implanted into the brain without the need for head-leveling or coordinate-mapping during surgery. The RatHat system can be used in conjunction with the traditional u-frame stereotaxic device, but does not require the use of a micromanipulator for successful implantations. Each RatHat system contains several primary components including the implant for mounting intracranial components, the surgical stencil for targeting drill sites, and the protective cap for impacts and debris. Each component serves a unique function and can be used together or separately. We demonstrate the feasibility of the RatHat system in four different proof-of-principle experiments: 1) a 3-pole cannula apparatus, 2) an optrode-electrode assembly, 3) a fixed-electrode array, and 4) a tetrode hyperdrive. Implants were successful, durable, and long-lasting (up to 9 months). RatHat print files are easily created, can be modified in CAD software for a variety of applications, and are easily shared, contributing to open science goals and replications. The RatHat system has been adapted to multiple experimental paradigms in our lab and should be a useful new way to conduct stereotaxic implant surgeries in rodents.

**Impact Statement:** We demonstrate a new approach to rodent stereotaxic surgery. Rodent neurosurgery is a complex skill that requires expensive equipment for head stabilization and micromanipulators for localization. The RatHat is a 3D printable brain implant system that reduces costs and time using pre-mapped and printed surgical files. A surgical stencil allows for quick placement of drill holes, and a RatHat places components in the brain using atlas coordinates. The RatHat system is an easily shared resource facilitating open science goals for simple replications and archiving of specific experimental applications.

## Introduction

Rodent neurosurgery is challenging to master, especially for implant surgeries involving multiple target sites. Long surgeries cause surgeon fatigue and distress in animals, which can affect recovery time and surgical outcomes (Ferry et al., 2014; Pritchett-Corning et al., 2011; Hoogstraten-Miller and Brown, 2008; Fox, et al., 2015). With an increased emphasis on circuit analysis that includes multiple brain targets (e.g. DREADDs and optogenetics), opportunities for positioning errors are increased (Jorgenson et al., 2015; Bassett & Sporns, 2017; Jayachandran et al., 2019).

Typically, brain implants are placed using a u-framed stereotaxic apparatus in which the rat’s head is stabilized with ear bars and a tooth bar, putting the rat into a three-dimensional atlas space (Paxinos and Watson, 2013). Micromanipulators attached to the u-frame allow implants to be precisely moved in xyz coordinate planes. However, this setup can introduce unrecoverable user errors that go unnoticed in the early stages. For example, while surgeons are trained to level the head in the anterior/posterior (A/P) plane, many fail to level in the medial/lateral (M/L) plane yielding asymmetrical implants/injections that produce time-consuming and expensive confounds unnecessarily increasing the number of animals needed for a study (Fomari et al., 2012; JoVE, 2019). Notably, there hasn’t been a major change in how neuroscientists approach stereotaxic methods in decades.

As a practical issue, a standard u-frame surgical apparatus can range from $5k-$50k (or more with addition of specialty add-ons), costing research labs a considerable portion of their equipment budgets and presenting a bar-to-entry for less well-funded laboratories.

Here we introduce a customizable, fully integrated 3D-printable stereotaxic brain implant system called RatHat that is freely available to academic researchers (Allen et al., 2017). The RatHat system can be used in conjunction with, or replace, the u-framed stereotaxic apparatus in neurosurgical methods requiring atlas-based positioning. A key feature is that the system self-aligns to atlas space because it fits the skull, eliminating the need for micromanipulator measurements and head-leveling.

The RatHat system reduces costs, training, and surgery time. It is customizable for a variety of surgical applications through modifications of the Computer Aided Design (CAD) environment prior to surgery (e.g., Autodesk, Blender, etc.). RatHat files are easily shared over the internet and archived for later use with versioning (aiding new experiments and replications). Printouts are considerably less expensive than similar commercial products while providing a larger range of implant possibilities. Furthermore, surgeons can map out coordinates in an interactive 3D environment to visualize the surgery prior to implantation, reducing demands during surgery.

RatHat applications have been adopted in our lab for a variety of experimental needs. Here, we demonstrate the use of RatHat in four experimental applications: multi-site chronic cannula, multi-site optrode-electrode combination implants, a fixed microwire microarray, and a tetrode hyperdrive with a microarray insertion tip.

RatHat is freely available to academic researchers, achieving open science goals. Academic researchers interested in receiving the 3D files can contact Dr. Timothy Allen. We will first provide you a license to be executed by your institution, and upon completion, 3D files of the implant system.

## Materials and Methods

RatHat components are printed using the 3D Systems ProJet 1200, a high resolution (56-micron xy, 30-micron layer thickness) 3D printer that uses micro-stereolithography (laser polymerization of resin and UV light-curing), but any high-resolution 3D printer can be used. With the ProJet 1200 prints, we use VisiJet FTX Green resin, a UV curable and biocompatible plastic composition commercially used in castings because it is a durable with a tensile strength of 30MPa (or 4,351 PSI). After devices are printed, we always ensure the holes are clear of debris or resin by thoroughly cleaning prints with multiple dips in a 70% isopropyl alcohol (in _di_H_2_0) solution and clearing holes by using a pressurized air output hose. Non-printable components such as wires or tubing are secured to the implant device prior to surgery with cyanoacrylate (Zap CA+, Super Glue Corporation, California) followed by a quick-cure spray (Zip Kicker, Super Glue Corporation, California). Another advantage of the RatHat system is that these components are easily assembled using build-specific 3D printable assembly bases or jigs. All implants are sterilized with 70% ethanol in diH_2_0 before surgical implantation and a gas sterilizer (ethylene oxide). Autoclaving is not recommended.

### *Components:* RatHat implant, Surgical Stencil, Protective Cap, and Implant Jig

Several components are common to all designs. The RatHat implant is a stable and secure housing apparatus for long-term neurosurgical implants. It is secured to the skull with anchor screws and dental cement. The underside of the RatHat that makes contact with the skull contains horizontal channels for dental cement, designed to optimize long-term adhesion of the implant to the skull and anchor screws (up to 9 months in our cohort). The version information and animal/experiment ID can be included on the print as well for ease of identification.

The surgical stencil contains all alignment and drill holes for the specific target sites needed in the surgery and was designed to facilitate rapid and accurate drilling of implantation and/or infusion sites to match the RatHat implant base. The surgical stencil is a transformative device for any surgeon to rapidly and cleanly introduce holes or craniotomies for an implant or injection. It is easy to print and uses relatively small amounts of resin, so multiple copies can be used for a single surgery in case a back-up is needed. This also helps with making straight and unbiased holes if free-handed drilling is preferred.

The protective cap safeguards other RatHat components (e.g. cannula tubes, dummies, electrodes, tetrode drives, etc.) from dust, debris, and impacts. It mounts on the side-walls of the RatHat implant base and is secured with a screw. The walls and the protective cap are outfitted with screw-holes for alignment on all sides to accommodate left- and right-handed surgeons. The protective cap can be printed with lab insignia and/or animal names for quick identification purposes. Protective caps can be replaced with a reprint if damaged in any way.

The jig (used to build cannula implants) serves to model the brain space and allows for precise placement and securing of implant components such as cannula tubes in the RatHat prior to surgery. In order to prepare the RatHat cannula implant base for surgery, the cured and cleaned 3D print is placed inside the jig. Next, pre-measured and cut stainless-steel tubes (27 ga, Component Supply Company, Sarasota, FL) are placed into the RatHat through the corresponding holes in the jig. The depths of the cannula are dictated by CAD-measured ledges printed within the jig for precise D/V depths. This reduces fabrication time and more importantly, measurement errors, as hand-cut cannula tubes do not need to be precision-measured to a discrete depth, since the jig dictates depth. Once the stainless-steel cannulae are secured to the RatHat implant base, the device can be implanted in the brain without the need for coordinate mapping during surgery. In this way, the jig replaces the dorsoventral (D/V) component of a stereotax micromanipulator arm, allowing for hand implants if the surgeon feels comfortable doing so.

### Animals and General Surgical Methods

Subjects used were Long Evans rats that weighed 250-275g on arrival (n=21, 2 female). All rats included were used for other primary experiments. Rats were individually housed in clear rectangular polycarbonate cages to ensure surgical implants were protected from damage by cage-mates. Rats were maintained on a 12hr light-dark cycle (lights off at 10:00 am). Naïve rats were briefly handled for 3 - 5 days after initial arrival. Access to food and water was unrestricted before surgery. All surgical and behavioral methods were in compliance with the Florida International University (FIU) Institutional Animal Care and Use Committee (IACUC) and Institutional Biosafety Committee (IBC).

Surgically implanting the RatHat follows basic techniques for intracranial survival surgery (refer to Fig. 1 for visualization). Briefly, general anesthesia was induced (5%) and maintained by isoflurane (1-2.5%) mixed with oxygen (800 ml/min). Rats were placed in a stereotaxic apparatus in the sterile surgical field for stabilization with ear- and tooth-bars (although RatHat surgery can be performed without this apparatus). Rats were administered glycopyrrulate (0.2 mg/ml, 0.5 mg/kg, s.c.) and 5 ml Ringer’s solution with 5% dextrose (s.c., over the duration of the surgery) for hydration. Temperature was monitored with a rectal thermometer and maintained within ±1C° of baseline temperature with a heating pad. The skull was exposed following a midline incision or fish-eye cut. The periosteum was detached from the skull using cotton-tipped applicators (Puritan Medical Products, Maine) and clamped with small hemostats to expose the width of the skull up to the lateral ridges (and 2 mm beyond the ridges when accessing more lateral structures) and 3-4mm length (A/P) beyond bregma and lambda. Score marks were made on the skull using the scalpel blade to aid dental cement adhesion.

**Figure 1:**
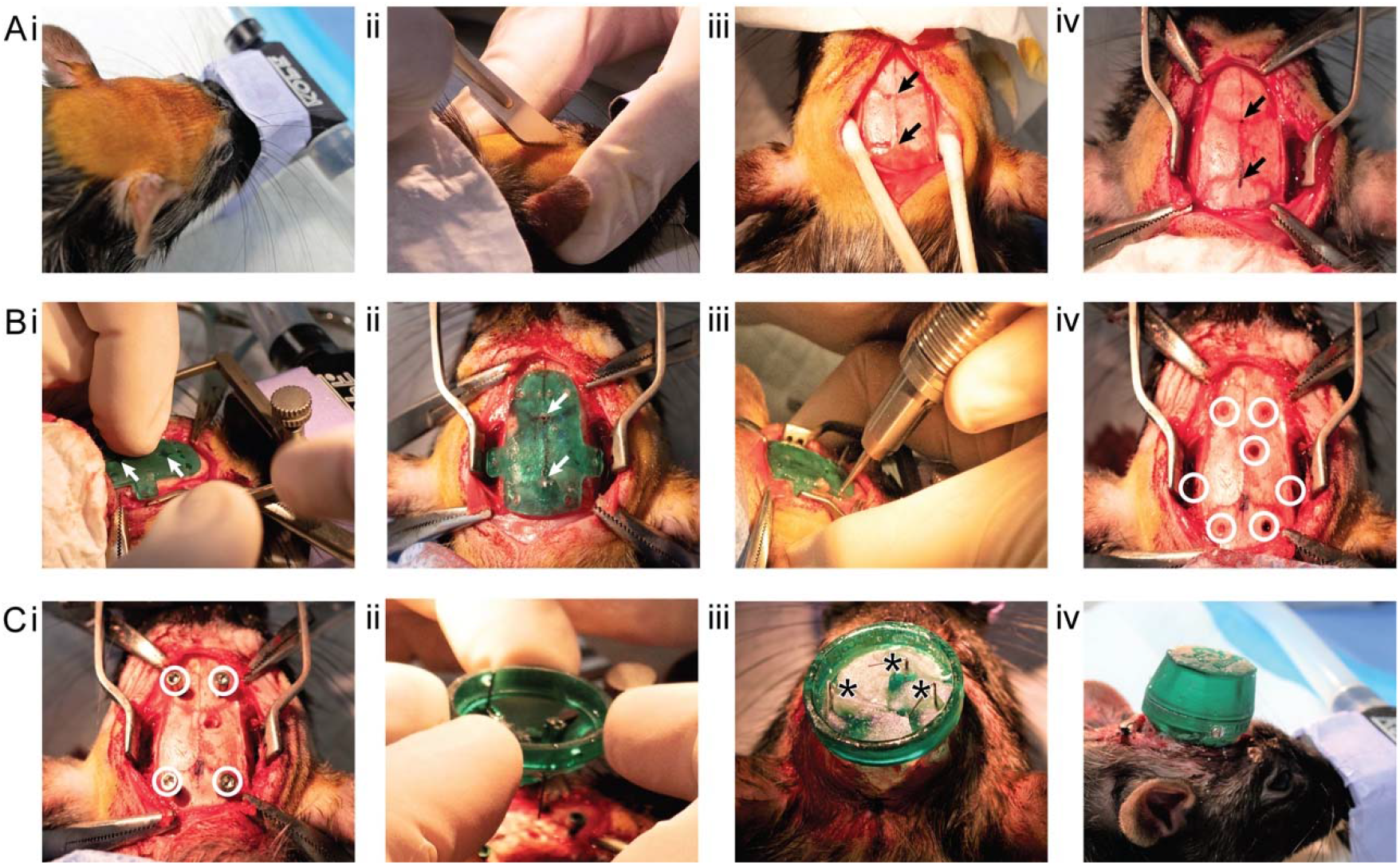
RatHat surgical procedures. **A) Prep: i.** Prepare the skin for incision. **ii.** Make the incision. **iii.** Clean skull and expose bregma and lamda. **iv.** Mark bregma and lamda and secure clamps to periosteum as needed. **B) Drill: i.** Place the stencil on the skull and align it to bregma and lamda. **ii.** Glue the stencil to the skull using cyanoacrylate and a quick-cure spray. **iii.** Drill holes according to the stencil. **iv.** Holes for skull screws and cannula (RE and PER) shown. **C) Implant: i.** Insert skull screws (holes remain for implant sites). **ii.** Manual placement of the preassembled cannula RatHat implant base. **iii.** Dental cement RatHat to the skull and insert dummies (asterisks indicate location of cannula poles). **iv.** Place the protective cap and secure with a screw.

The surgical stencil was aligned to bregma and lambda using the landmark holes that are surrounded by crosshairs to facilitate visualization and placement. The stencil was secured to the skull using cyanoacrylate (Zap CA+, Super Glue Corporation, California) followed by a quick-cure spray (Zip Kicker, Super Glue Corporation, California). Drill holes were made in the appropriate regions according to the specific RatHat build using a surgical drill (OmniDrill 3S, World Precision Instruments). Dura mater was ruptured at the implant sites using a 32-gauge needle. The stencil was removed using a scalpel blade or a spatula and discarded. Excess cyanoacrylate residue was scraped off the skull to clear any debris that could interfere with placement of the RatHat implant base. The skull was thoroughly cleaned with sterile saline or hydrogen peroxide (avoiding contact with skin and muscles) to ensure successful long-term adherence of the RatHat. Titanium anchor screws were secured into place. The RatHat was aligned to the drill holes and carefully lowered into place using the micromanipulator arm or by hand, fitting it flush with the skull. Dental cement was applied in layers to secure the base to the anchor screws and skull using a wooden applicator tip, a syringe, or a paint brush (saturated first in the curing liquid and then used to pick up the dry powder, which polymerized into the cement and facilitated creation of a smooth and well-anchored implant, free of jagged edges). The inside of the implant was filled with dental cement to further stabilize components. Once dry, the protective cap was secured onto the wall of the RatHat using a small screw. The posterior incision was sutured if necessary, rats were administered an analgesic (Flunixin, 50 mg/ml, 2.5 mg/kg, s.c.), and topical antibiotic ointment was applied around the surgical incision. The rat was placed in a post-surgical recovery incubator until awake and moving, and then returned to a clean home cage. A day following surgery, rats were given an analgesic (Flunixin, 50 mg/ml, 2.5 mg/kg, s.c.) and topical antibiotic ointment was applied. The protective cap was removed to check that the RatHat implant components were in good condition. Rats were monitored post-operatively for a week and then resumed behavioral or experimental testing.

Upon completion of the experiments, intracranial placements were mapped using postmortem brain slices. Briefly, rats were induced under general anesthesia using isoflurane (5%) and transcardially perfused with 100 ml of ice cold 0.1M PBS followed by 200 ml of 4% paraformaldehyde (pH 7.4; Sigma-Aldrich, St. Louis, MO). Brains were post-fixed overnight in 4% paraformaldehyde and then cryoprotected in a 30% sucrose and 0.1M PBS solution prior to sectioning (Leica CM3050S, Leica Biosystems). Three sets of immediately adjacent sections (40 μm, coronal orientation) were saved. One set was mounted onto microscope slides for a cell-body specific Cresyl Violet stain for placement analysis.

## Results

***Experiment 1:* 3-pole cannula RatHat for simultaneous implantation of multiple cannula (Fig. 2)**. Commercially available multisite cannula assemblies from vendors such as PlasticsOne and WPI are custom ordered, requiring a necessary lead-time, and very expensive. Furthermore, they only accommodate up to two cannulas anchored together by a thin plastic tether, and are unable to incorporate poles for angled insertions (other than perpendicular to the skull).

**Figure 2:**
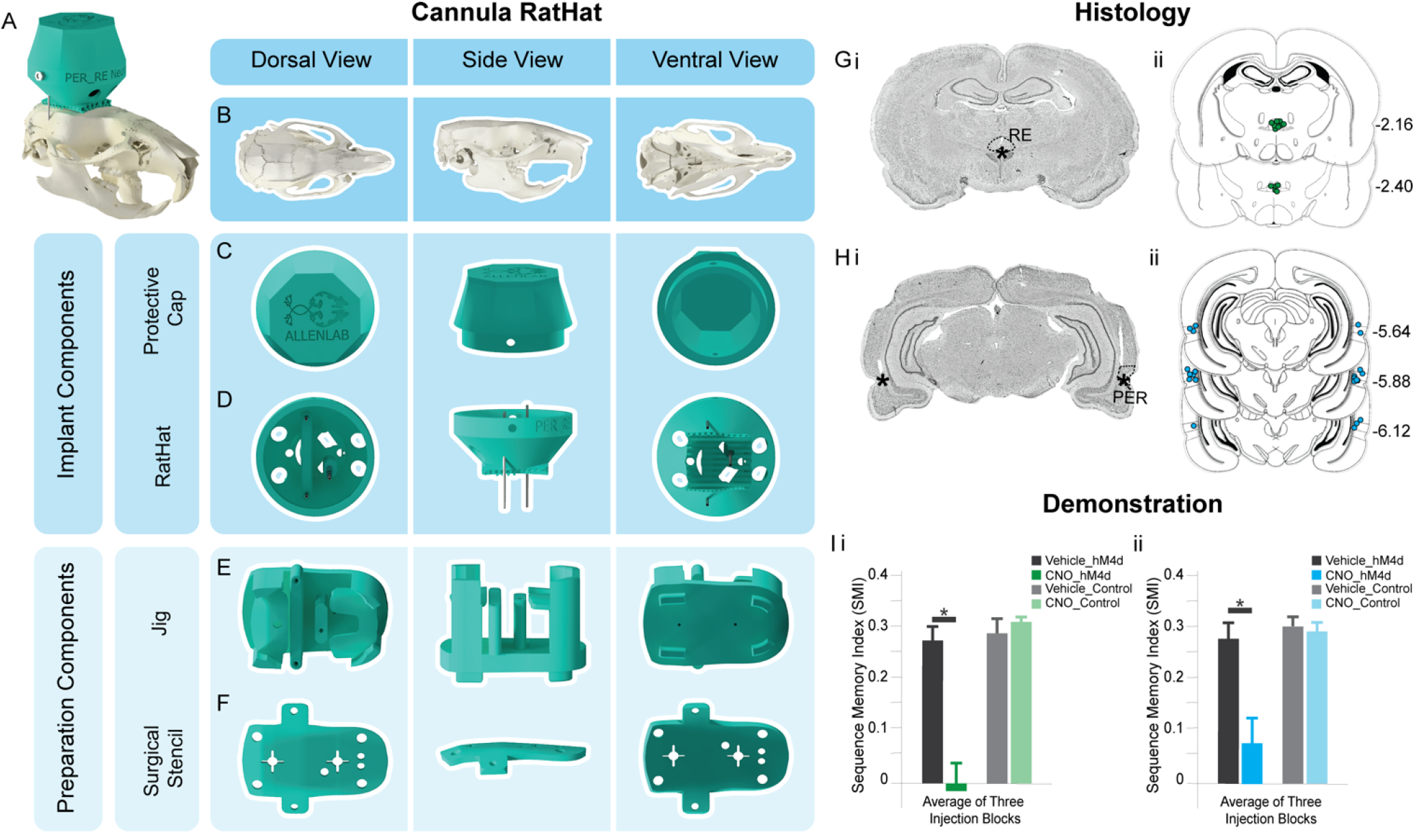
3-Pole Cannula RatHat. **A)** Full RatHat cannula assembly on the skull. **B)** Rat skull in different orientations. **C)** The Protective Cap shown in different orientations. **D)** The RatHat Implant Base with preassembled cannula. **E)** The Jig is used to assemble the cannula tubes into the RatHat implant prior to surgery. **F)** The stencil contains reference marks for bregma and lamda to align to the skull; once adhered, all drill marks are properly placed for cannula access points and anchor screws, saving time. The stencil is removed and discarded after drill holes are made. **G)** i. Sample coronal slice. The asterisk indicates the infusion cannula tip location in RE. **ii.** Microinfusion injector tip location in the RE for all rats (n=13). Numbers to the right of each section indicate distance (mm) anterior to bregma. **H) i.** Sample coronal slice. The asterisks indicate the infusion cannula tip location in PER. **ii.** Microinfusion injector tip location in the PER for all rats (n=13). Numbers to the right of each section indicate distance (mm) anterior to bregma. **I)** Rats were injected with AAV-hM4Di (an inhibitory DREADD) in mPFC or a control virus, and a cannula targeted RE and PER (bilaterally). Well-trained rats were infused with CNO in RE and PER (the DREADD agonist) or vehicle prior to testing. **i.** Silencing the mPFC → RE terminals (the CNO-hM4Di group) abolished sequence memory. **ii.** Silencing the mPFC → PER terminals (the CNO-hM4Di group) abolished sequence memory. G-I were reprinted from Jayachandran et al. (2019) with permission. *Abbreviations:* (CNO) clozapine N-oxide, (mPFC) medial prefrontal cortex, (RE) nucleus reuniens of the thalamus, (PER) perirhinal cortex.

The RatHat cannula system contains multiple pre-measured cannulas assembled before surgery, reducing surgical time by eliminating the need to identify coordinates with micromanipulators and make insertions one-at-a-time. Here, two cannulas targeted perirhinal cortex (PER) bilaterally, and one cannula targeted the nucleus reuniens of the thalamus (RE). PER is a good site to demonstrate the RatHat cannula approach because it is a difficult structure to access, given its depth and laterality (A/P −3.0 to −7.0; M/L ±.7.2; D/V −6.5 to −7.5; Paxinos & Watson, 2013; Burwell, 2001). The third cannula targeted RE, a structure that lies directly below the superior sagittal sinus (SSS; A/P −1.08 to −3.48; M/L ±.08; −6.8 to −7.8 D/V). The SSS can easily rupture, prolonging surgical time and causing significant damage or death. Thus, we incorporated an angled cannula pole (10°) into this RatHat design to target RE and avoid SSS. This angled pole is fitted with a depth-stop, eliminating the need for D/V measurements, and was inserted by hand then secured to the RatHat implant base.

Prior to implantation, male rats (n=13) were trained in an odor sequence memory task (from Jayachandran, et al., 2019). Briefly, once rats reached criterion in the task, they underwent RatHat implantation surgery. Following recovery, rats were retrained on the sequence task until they reached performance criteria. This task demonstrates the durability of the RatHat, which is an ideal device for experiments that require extended testing periods and involve extensive task related wear-and-tear. These rats completed approximately 60 sessions after surgery, with 200-300 nose-pokes/session. Additionally, rats were given 12 infusions over several weeks to either PER (bilaterally) or thalamic RE. Infusions targeted the structures of interest and resulted in distinct sequence memory disruptions that relate to the functioning of those regions. RatHat implants stayed on for an average of 6 months (maximally 9 months), when rats were killed for histological analysis. There were no significant deviations from the targets (see Fig. 2 G/H) showing great reliability.

***Experiment 2:* RatHat design for a combination of optogenetics and microwire recordings (Fig. 3)**. We implanted rats (n = 4; 2 females) weighing approximately 275-350g at surgery. Here, the optrode targeted the junction of the thalamic RE body and thalamic RE wing (perireuniens; −2.3 A/P, - 0.5 M/L, −7.0 D/V) and was implanted vertically at an M/L slightly lateral to the midline (a different approach compared to the cannula; Viena, et al., 2019). The surgical stencil for this version was designed with drill hole guides for the injection site/optrode, stainless steel wire electrode, and anchor screws. After the holes were drilled, an injection of AAVr-CAG-hChR2-H134R-tdTomato (experimental virus to express channelrhodopsin; Addgene cat #: 28017) or pAAVr-CAG-tdTomato (control virus; Addgene cat #: 59462) was made using pulled glass pipettes (P-2000 Laser-Based Micropipette Puller, Sutter Instruments) with a tip diameter between 80-100 μm driven by a motorized infusion pump (0.3-0.5 μL at 60 nL/min; Nanoject III, Drummund Scientific). Because the optrode has a built-in depth-stop for the D/V axis, a jig was not required for this version. Similar steps for implantation were used during this surgery as described above.

**Figure 3:**
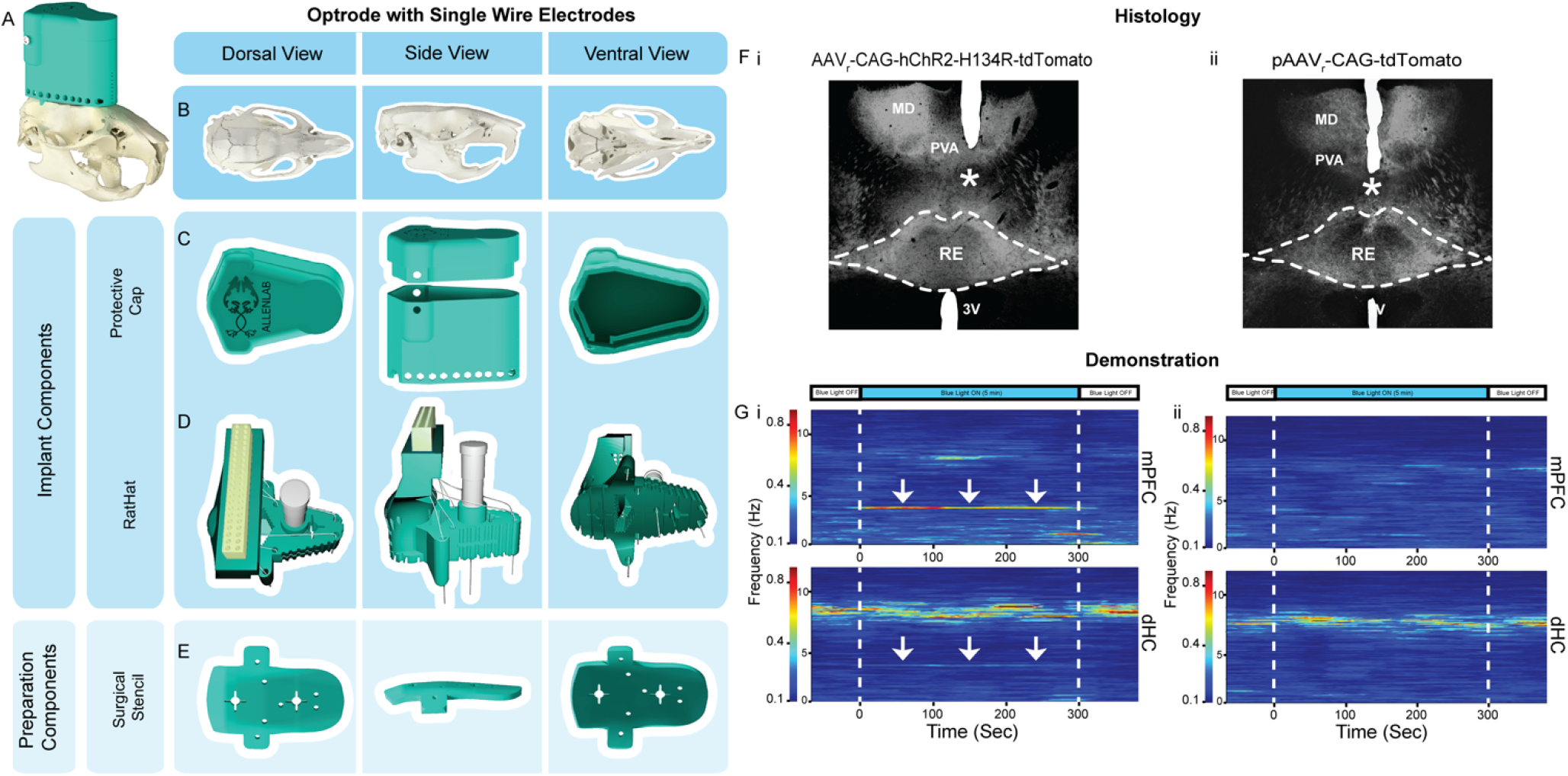
Optrode and SS Wire Electrode RatHat Assembly. **A)** Optrode/Electrode RatHat implant on an average sized rat skull. **B)** Rat skull viewed across different anatomical planes. **C)** View of the RatHat protective cap and wall in different orientations. These items protect the internal components post implantation. **D)** RatHat implant base (in green) preassembled with optrode and electrode single wires. **E)** RatHat surgical stencil showing prefabricated holes that correspond to brain coordinates of interest, bregma and lamda, and screw locations for rapid drilling on the skull. **F)** Brain sections showing channelrhodopsin (ii) and viral control (ii) expressed neurons in the midline thalamus and the optrode placement (asterisk) just above RE in representative cases, demonstrating the effectiveness of using the RatHat system. **G)** Perievent spectrograms of representative mPFC and dHC LFP showing the 5 minute period in which the blue LED light was administered via the optrode (see asterisk for tip location) activating ChR2 ion channels in infected (i; AAVr-CAG-hChR2-H134R-tdTomato) and control (ii; pAAVr-CAG-tdTomato) rats. Also shown, 60 seconds before and after the stimulation block. Pulsed blue light activation (4Hz, 60 ms pulse width) of RE ChR2+ neurons elicited a 4Hz frequency rhythm (see arrows) in the mPFC (strong) and dHC (weak) LFP signal. We also observed comparable frequency-specific activations at 1Hz, 2Hz, and 8 Hz. This change, however, was not observed in control animals (on right). *Abbreviations:* (mPFC) medial prefrontal cortex, (dHC) dorsal hippocampus, (RE) nucleus reuniens of the thalamus, (PVA) paraventricular nucleus, (MD) medial dorsal nucleus, (ChR2) channelrhodopsin.

We show sample data in which optogenetic stimulation of RE in experimental rats yielded a 4Hz frequency rhythm in the mPFC (strong) and dHC (weak) LFP signal, but not in controls, demonstrating efficacy of the optogenetic approach (Fig 3G). Implants remained in place for 4.5 months until brain analysis. Proper placement of the optrode and electrode wires was verified and consistent in all rats (see Fig. 3F for representative slices depicting optrode placements).

***Experiment 3:* RatHat for implanting fixed stainless-steel wire arrays** targeting prelimbic (PL) and infralimbic (IL) regions of the medial prefrontal cortex (mPFC; see Fig. 4) were piloted for feasibility in male rats (n=2; ~350g at surgery). The electrode array was built similar to those used in other experiments (Krupa et al., 2009; Narayanan et al., 2006). The surgical stencil for this version included a craniotomy window supporting the electrode arrays bilaterally targeting PL/IL (1.7-4.0 A/P, ±1.8 M/L, −3.0 D/V) in addition to landmark and anchor screw holes. Once the craniotomy and drill holes were made, anchor screws were inserted. Next, the RatHat implant base was secured to the skull. The electrode array was then inserted into the brain, docked into place on the RatHat base, and secured with dental cement. The protective wall was then secured with dental cement (filling in the base up to the Electrode Interface Board). Once dry, we plugged the rat into the electrophysiological recording system (Plexon, Dallas, TX) to assess neural activity. After, the protective cap was placed and secured to the RatHat. Neural activity was assessed over the course of the next several months. RatHat electrode arrays remained in place for approximately 4 months with good signal. We successfully recorded well-isolated single-unit activity (two well-isolated units are shown in Fig. 4H). Marking lesions were performed using a NanoZ for localizing electrode sites in the brain (Fig. 4G).

**Figure 4:**
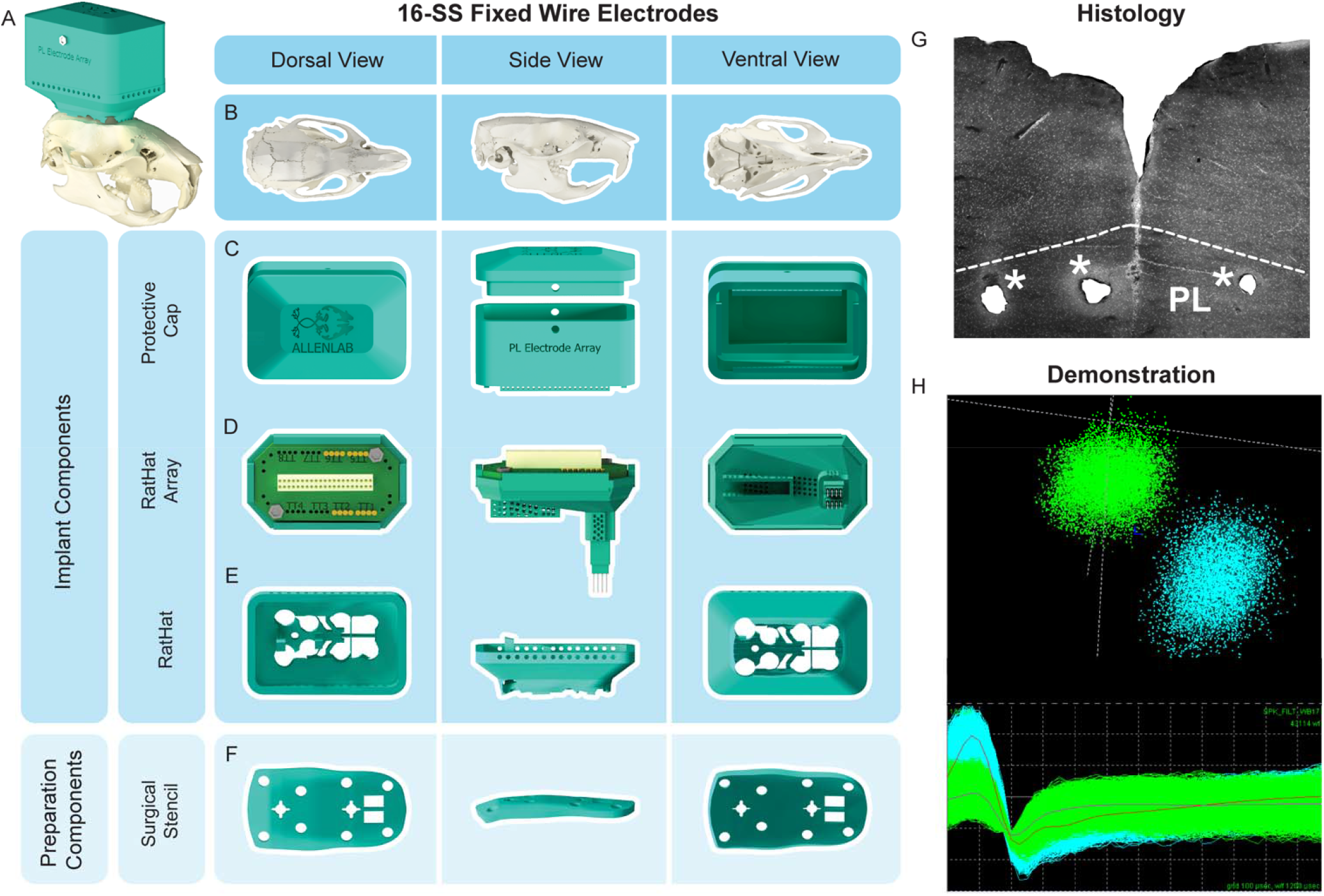
16 Single Wire Fixed Electrode Array RatHat. **A)** Fixed electrode array RatHat on the skull. **B)** Rat skull in different orientations. **C)** The protective cap and wall that protects the electrode array after implantation. **D)** The RatHat electrode array with preassembled fixed stainless-steel single wires docks into the RatHat implant base. **E)** The RatHat implant base is anchored to the skull before the RatHat electrode array is docked, ensuring the wires descend to the correct DV. **F)** The Stencil contains reference marks to align bregma, as well as a craniotomy window and drill holes for anchor and ground screws. **G)** Sample coronal slice of the 16-Wire SS Electrode Array RatHat. The asterisks indicate single wire tip locations. **H)** Sample cluster plot showing two isolated mPFC units on a single channel during free-roaming behavior. *Abbreviations:* (PL) prelimbic cortex.

***Experiment 4:* 8-tetrode hyperdrive RatHat** (Fig. 5). Microdrive screws and shuttles for the RatHat hyperdrive were assembled and implanted similar to others (Wilson and McNaughton, 1993; Gray, et al., 1995; Nguyen, et al. 2009). The tetrode array was securely encased in the protective wall prior to surgery. We implanted the RatHat hyperdrive in male rats (n=2; ~350g at surgery). This stencil version was the same as that used in Experiment 3. After the RatHat implant base was secured to the skull, the RatHat hyperdrive was carefully placed by hand and secured onto the base with dental cement. Immediately after, tetrodes were driven 1 mm and the rat was plugged into the electrophysiological recording system (Plexon, Dallas, TX) to assess signal on the wires. Once functionality was established, the rat was unplugged and protective cap was secured into place. Tetrodes were driven 250μm/day until reaching a depth of 2.8 to 3.0 mm (staggered) with a goal of recording from mPFC cells (4.7 to 2.5 A/P range, ±0.2 to ±1.6 M/L range). The hyperdrive successfully isolated single-units in mPFC of freely-behaving rats, demonstrating the RatHat application (Fig. 5H). Four weeks after implantation, marking lesions were performed using the a NanoZ to localize tetrode sites. Cresyl Violet-stained sections were analyzed for placement of the tetrode wires (Fig. 5G).

**Figure 5:**
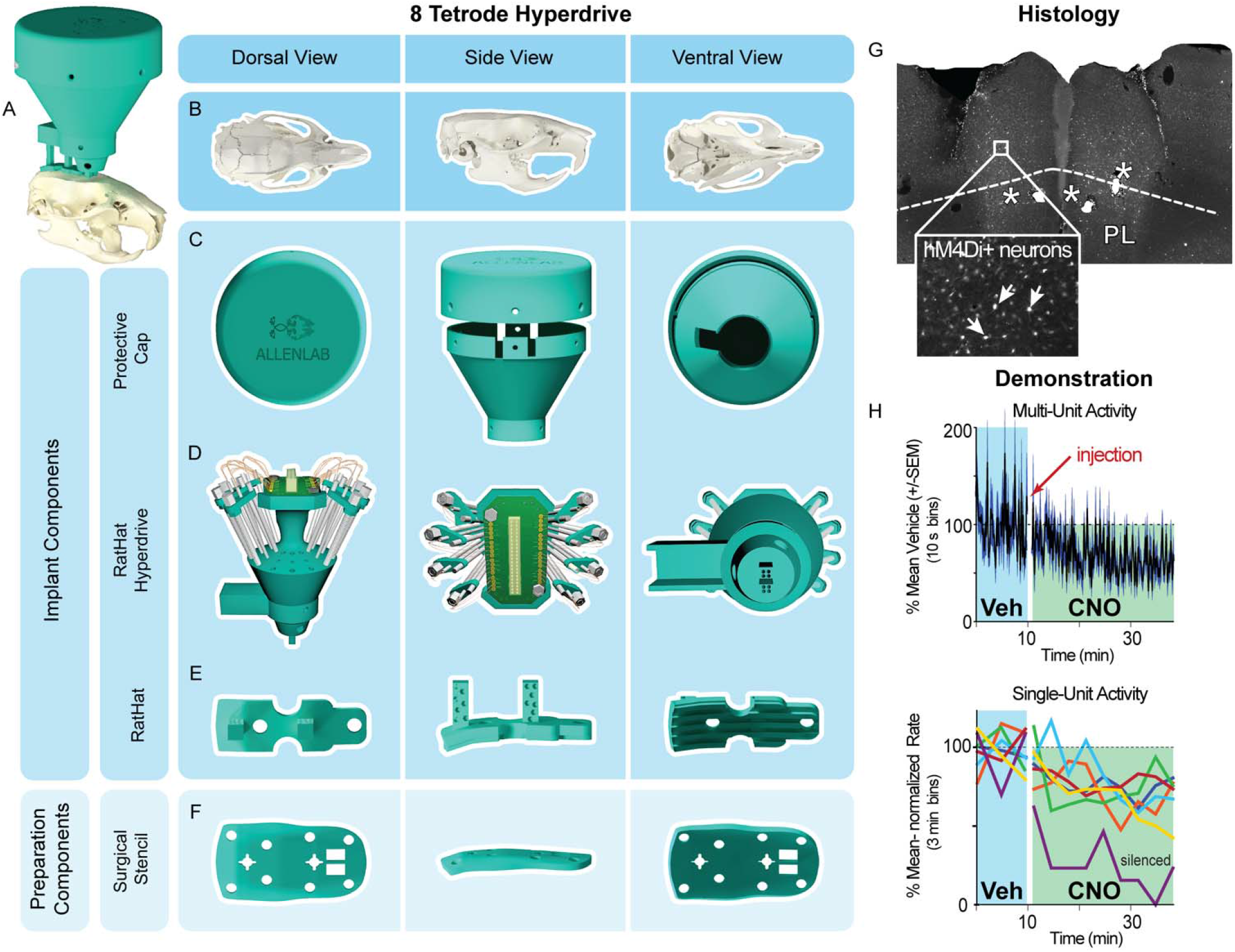
8-Wire Tetrode Hyperdrive RatHat. **A)** The fully assembled 8-Wire Tetrode Hyperdrive RatHat on the skull. **B)** Rat skull in different orientations. **C)** The protective cap and wall ensure the RatHat hyperdrive is safe from impacts and debris. **D)** The hyperdrive with preassembled drivable tetrodes targeting regions in mPFC. **E)** The RatHat implant base is secured to the skull and has docking poles on which the RatHat hyperdrive sits, ensuring the tetrode tips are placed right above cortex. **F)** The stencil aligns to bregma and lambda and contains guide holes for drilling craniotomies and anchor screw holes. **G)** Sample slice with 8-wire tetrode hyperdrive RatHat. Asterisks indicate the tetrode wire tips **H)** Implanted tetrodes in mPFC with hM4Di expression showing functional inhibition following CNO injection (1mg/kg). *Abbreviations:* (CNO) clozapine N-oxide, (mPFC) medial prefrontal cortex, (PL) prelimbic cortex, (Veh) Vehicle.

## Discussion

RatHat is a 3D-printed stereotaxic device that can be used for a range of applications, such as cannula placements, microinfusions, optogenetics, and electrophysiological recordings. The RatHat system was developed to improve surgical accuracy and precision, reduce surgery time in rodent neurosurgical procedures, and contribute to open science goals. The RatHat system is freely-available to academic researchers. This is a major change to current stereotaxic approaches because we replaced an approach that has been used for several decades that uses micromanipulators for measurements during surgery. The RatHat system saves time, money, offers reliability, and provides for surgical replications. The fundamental system consists of complementary components including a RatHat implant base, a surgical stencil, and a protective cap. Here we demonstrated four different RatHat systems for feasibility in multiple types of neuroscience experiments. We verified the durability of these implants, which remained in place for up to 9 months, in spite of movement- and impact-dense behavioral tasks (e.g. Jayachandran, et al., 2019).

We plan to develop a RatHat for other commonly used species in neuroscience, including mice. We have also developed a version for chronic implants in the domestic pig, which facilitates surgery without the need for a traditional large-animal stereotaxic apparatus (HogHat; US Patent 10,251,722 B1, 2019). In addition, RatHat versions for other common neurosurgical applications are underway including an acute implant device that allows for single injections of excitotoxins, AAVs, DREADDs, etc.

Again, we make the RatHat freely-available for all academic researchers to aid in their experiments and contribute to open science goals. We look forward to new builds and implementations from other academic research groups.

## Acknowledgements

We thank Jason Hays for helping with figures, and Chrystle and Catherine Cu (CocoFloss) for donating UV-light curing dental cement.

